# Sexually divergent effects of social dominance on chronic stress outcomes in mice

**DOI:** 10.1101/2020.02.04.933481

**Authors:** Stoyo Karamihalev, Elena Brivio, Cornelia Flachskamm, Rainer Stoffel, Mathias V. Schmidt, Alon Chen

**Affiliations:** Department of Stress Neurobiology and Neurogenetics, Max Planck Institute of Psychiatry, Munich, Germany; International Max Planck Research School for Translational Psychiatry (IMPRS-TP), Munich, Germany; Department of Neurobiology, Weizmann Institute of Science, Rehovot, Israel

**Keywords:** Sex differences, stress, social behavior, social dominance, chronic mild stress, principal component analysis

## Abstract

**Background:** Sex and social context are two major factors in the development of depression and other stress-related disorders. However, few studies of the effects of stress on rodent behavior and physiology have investigated social context and fewer still have assessed the possibility of sex-specific effects of social context.

**Methods:** We assessed social dominance of group-living mice during several days of monitoring using a high-throughput automated behavioral tracking system. We then exposed groups from each sex to a three-week chronic mild stress (CMS) procedure, followed by a behavioral test battery. Finally, we used principle component analysis and post-hoc tests to explore the sources of variance in the behavioral outcome data.

**Results:** We found stable hierarchies in both sexes, however social dominance in males exhibited several additional associations with behaviors related to locomotion and exploration that were not seen in females. Crucially, pre-stress social dominance status was associated with opposing outcomes on multiple behavioral readouts between the two sexes following CMS. In particular, subordinate male mice and dominant female mice appeared more responsive to the environmental challenge, as observed in anxiety-like and locomotor behaviors.

**Conclusions:** This work demonstrates that sex differences interact with preexisting social dominance status to alter the effects of chronic stress. It highlights the importance of understanding the interplay between sex and social context and its contribution to individual differences in stress response.

## Introduction

Stress-related psychopathologies, such as mood and anxiety disorders, show a pronounced gender bias in their prevalence, severity, age of onset, and most common comorbidities(1–4). For example, the prevalence of major depressive disorder among women is two to three times higher than in men and is characterized by increased symptom severity(5,6). In women, major depression is more often comorbid with anxiety disorders, eating disorders, and sleep disturbances, while men with major depression are more prone to develop aggression, alcohol or substance abuse, and suicidal ideation(6,7).

Given these differences and the known sex dimorphism in stress responses in humans(8), it is surprising that few studies in rodent models consider the effects of sex when characterizing models of stress-related psychopathologies(9,10). Symptomatology of stress-related pathologies span several domains functioning including mood, metabolism and sociability. Importantly, differences in social behavior and social cognition prior to disorder onset may contribute to disorder susceptibility, especially considering that abnormalities in social functioning are an essential part of the symptomatology of stress-related disorders. Nonetheless, very few studies have looked into pre-existing differences in social behavior between the sexes and how those might contribute to the sexually divergent consequences of stress exposure.

Here, we explore the impact of social context in shaping the response to chronic stress. In particular, we investigate social dominance, a well-studied and central feature of rodent social groups. Wild and laboratory rodents form complex and dynamic social structures which typically involve the formation of dominance hierarchies(11) to improve social stability and reduce severe conflicts and aggression(12). As a consequence, an individual’s position in the dominance hierarchy has important consequences, including preferential access to food, shelter, and mates(13). Social rank within male hierarchies is also known to influence health, hormonal profile, brain function, metabolism and mortality(14,15). For instance, subordinate individuals display increased anxiety-like behavior, a suppressed immune response, higher basal corticosterone levels, and reduced life span(16). These types of relationships have classically been studied in male animals, as female mice have usually appeared more communal and displayed limited aggression(17). Recent work, however, has demonstrated that female laboratory mice also form hierarchies that appear quite similar to those seen in males, accompanied by some of the same dominance-related physiological markers, such as differences in corticosterone levels(18–22). Thus, we examined social dominance status as a putative mediator of sex differences in the response to adverse events.

To do so, we took advantage of a high-throughput automated behavioral monitoring system (Social Box, SB) to assess and better understand the hierarchies of groups of male or female mice(23,24). We then exposed mice to an established chronic stress procedure, the chronic mild stress (CMS) paradigm, and evaluated its effects using a series of standard behavioral and physiological readouts. Finally, we used baseline social dominance status to predict CMS behavioral outcomes. We hypothesized that an individual’s standing in the social hierarchy would be a potent predictor of behavior upon stress exposure, and that this relationship may differ between the sexes.

## Results

### Male and female dominance hierarchies

We first explored the hierarchical structure of grouped CD-1 mice over four days of baseline monitoring as well as the stability of hierarchies following an acute stressor (15 min of restraint stress). Social dominance was assessed by calculating groupwise normalized David’s Scores (DS), a well-established method for inferring social hierarchies(25,26). We based the DS on the numbers and directionality of chases between each pair of individuals in a group. A cumulative DS for the four baseline days of the SB assessment was used as a final measure of social dominance. In line with previous studies(18–22), we were able to detect a stable hierarchical structure in both sexes (**Figure 1a**).

**Figure 1.**
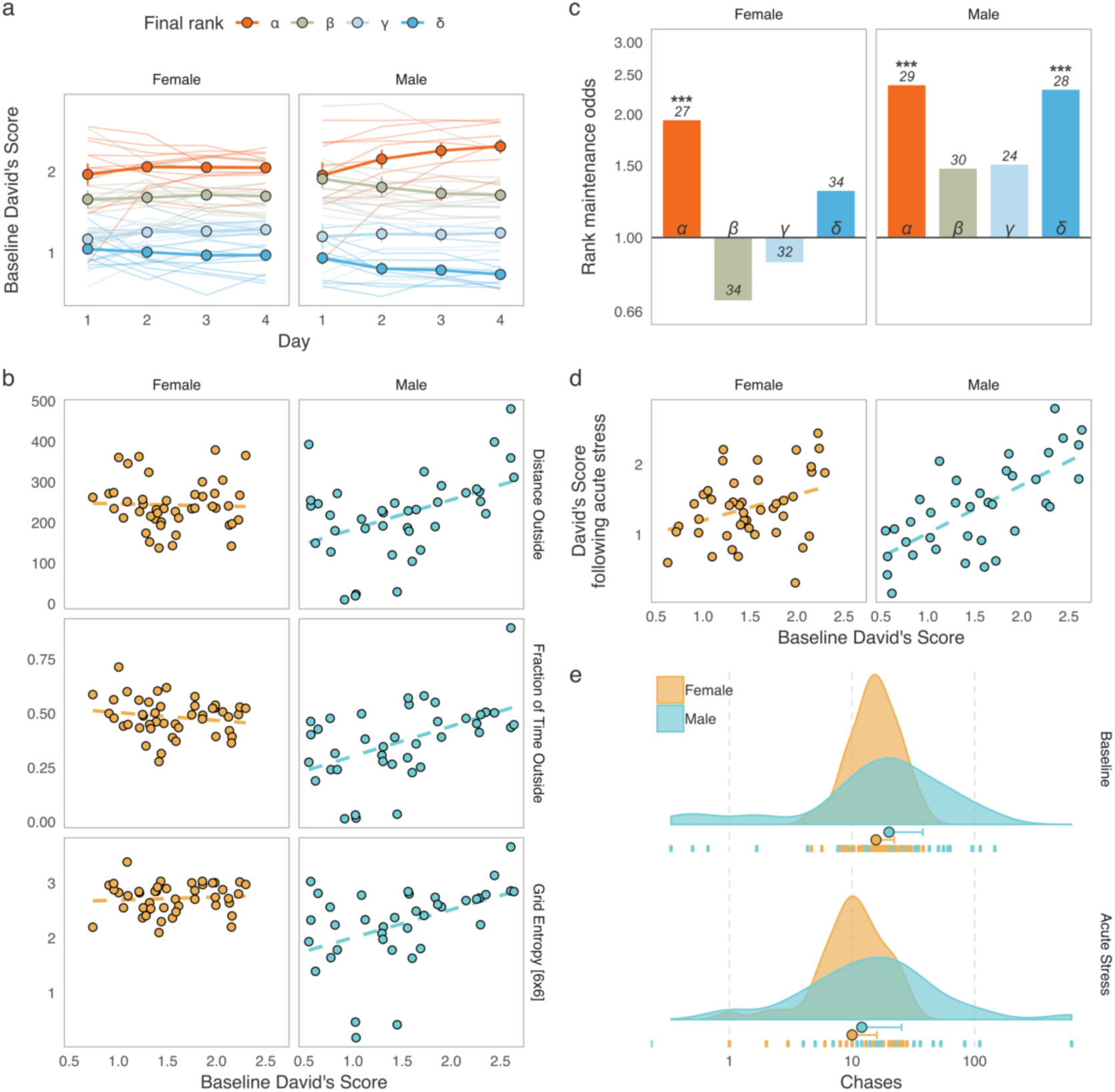
Social dominance hierarchies in males and females. (a) David’s Scores (DS) based on chases during the four baseline days of Social Box (SB) assessment show relatively stable social hierarchies in both male and female groups (each line represents an individual, colors represent the cumulative social rank on day 4, points are mean values for each rank ± standard error of the mean). (b) Male-specific associations between social dominance scores and behaviors related to locomotion and exploration. Dominant males had increased overall locomotion, spend more time outside the nest, and moved through the SB environment in a more unpredictable manner. These associations were not found in females. (c) Rank maintenance odds over the four-day baseline period. Depicted are odds of maintaining the same rank between consecutive days relative to chance-level (25%). Data is summarized according to the cumulative social rank on day 4; numbers indicate the number of individuals per rank. (d) Baseline DS predicts DS following acute restraint stress in both sexes, indicating that social dominance hierarchies may be relatively robust against acute stress. (e) Numbers of chases in male and female groups at baseline as well as following acute restraint stress. Both sexes display significantly fewer chases during the dark phase following an acute physiological stressor.

We further calculated several properties of male and female hierarchies, including steepness, despotism, directional consistency, and Landau’s modified *h*’, a measure of hierarchy linearity(27,28) (Supplementary **Figure 1**). Male hierarchies were more linear, more despotic, and had higher directional consistency than those of females. Interestingly, mice housed in larger groups show analogous relationships between sexes(19).

To investigate if social rank carries any sex-specific implications for overall behavior, we analyzed the correlation structure between an individual’s DS and 60 behavioral readouts recorded post-habituation (days 2-4) in the SB within each sex (Spearman’s rank correlation; for a detailed list of behaviors and how they are computed, see ref.(24)). Thirty-six of the behaviors tested (60%) showed significant correlations with cumulative baseline DS in at least one of the sexes (Supplementary **Figure 2**, *q* < .1, Benjamini-Hochberg adjustment within each sex). While the overall association pattern was quite similar between male and female mice, there were several strong correlations seen in males that were absent in females (**Figure 1b** and **Supplementary Figure 2**). These included Distance Outside – a measure of overall locomotion (Male *r*_*s*_ = .43, *n* = 40 mice, *p* = .006. Female *r*_*s*_ = -.09, *n* = 48 mice, *p* = .55), Fraction of Time Outside – the mean proportion of time a mouse spent outside the nest (Male *r*_*s*_ = .498, *n* = 40 mice, *p* = .0014. Female *r*_*s*_ = -.159, *n* = 48 mice, *p* = .29), and two related measures of roaming entropy, which assess the predictability of how an individual explores their environment – Entropy and Grid Entropy [6×6] (for brevity we only report the latter, Male *r*_*s*_ = .482, *n* = 40 mice, *p* = .0024. Female *r*_*s*_ = .097, *n* = 48 mice, *p* = .52). Correlations in males that were not present in females indicate that unlike in males, in females overall locomotion and exploration of the home environment are largely independent of social rank. Interestingly, no correlations were present in females but absent in males. Altogether, these findings suggest that male and female social dominance hierarchies, despite having a similar structure, may carry somewhat different consequences for overall behavior.

Next, we estimated DS stability over time by examining the frequency of rank change events and comparing those to the chance-level expectation. Briefly, normalized daily DS values were ranked for each group to create a four-rank hierarchy: *α* (dominant), *β, γ, δ* (subordinate) and each mouse was assigned a single rank based on its four-day cumulative chase DS. For each pair of consecutive days, we observed how many individuals maintained the same rank they had been assigned on the previous day.

We then calculated the rank maintenance odds for animals in each final rank category relative to the expected chance level (**Figure 1c**). The true probability of rank maintenance in our data was higher than chance in *α*-females, *α*-males, and *δ*-males (one-tailed binomial tests against the rank maintenance probability of 25%, *α*-females: 13/27 successes, *p* = .008, *α*-males: 17/29 successes, *p* = 12×10^−5^, *δ*-males: 16/28 successes, *p* = 3×10^−4^). These results indicate that the highest rank in a hierarchy is often occupied by the same individual over time in both sexes, while the lowest rank appeared to be stable in males only (**Figure 1c**).

In addition to stability over time during baseline recordings, individual DS also remained relatively stable following acute restraint stress (Pearson’s correlation between cumulative baseline DS and DS following acute restraint on day 5, **Figure 1d** and Supplementary **Figure 1**). Both males (*r* = .699, *n* = 37 mice, *p* = 1.46 × 10^−6^) and females (*r* = .338, *n* = 44 mice, *p* = .025) showed significant DS correlations from baseline to acute restraint. Finally, we investigated the possibility of differential effects of acute restraint on the behavior used to produce the DS – numbers of chase events (**Figure 1e**). Repeated-measures ANOVA on log-transformed chase numbers showed that the number of chases decreased significantly following acute restraint stress (*F*(1, 78) = 30.559, *n* = 84 mice, *p* = 4.15×10^−7^), however the extent of this decrease did not differ between the sexes (Sex x Stage interaction, *F*(1, 78) = .023, *n* = 84 mice, *p* = .881).

The apparent robustness of social hierarchies over time and in response to acute stress suggested that predictions from the baseline assessment may carry information that would still be relevant to behavioral outcomes following a long-term intervention. More specifically, we hypothesized that occupancy of the highest-ranking positions in the social hierarchy in both sexes and additionally the lowest in males might be sufficiently stable to allow for long-term predictions.

### Effects of chronic mild stress on behavior and physiology

To investigate the effects of pre-existing social dominance status on the behavioral response to chronic stress, we employed a CMS protocol adapted for group-housed animals.

In short, groups were exposed to a weekly schedule of two daily randomly combined mild stressors (e.g., wet bedding, tilted cage, overcrowding) for a total of three weeks. Six groups of each sex (*n* = 24 per sex) were randomly assigned to receive CMS, while the rest of the groups (six groups of females and four groups of males) were assigned to the control condition. The 21-day CMS procedure was followed by a behavioral test battery for both control and CMS mice, which included tests previously shown to capture the effects of chronic stress (**Figure 2a**). This included, among others, classical tests of locomotion (open field test, OFT), anhedonia (sucrose preference test), anxiety-like behavior (elevated plus maze, EPM), and stress coping (tail suspension test). Additionally, we assessed several physiological indicators of stress level (**Figure 2b-e**). All the physiological and behavioral outcome variables following CMS were collected into a single dataset (**Figure 2f**). Since the full experiment was run in two batches, all outcome variables were adjusted for batch effect (*see* Methods). To improve readability, we report the batch-adjusted values relative to the mean of female control mice.

**Figure 2.**
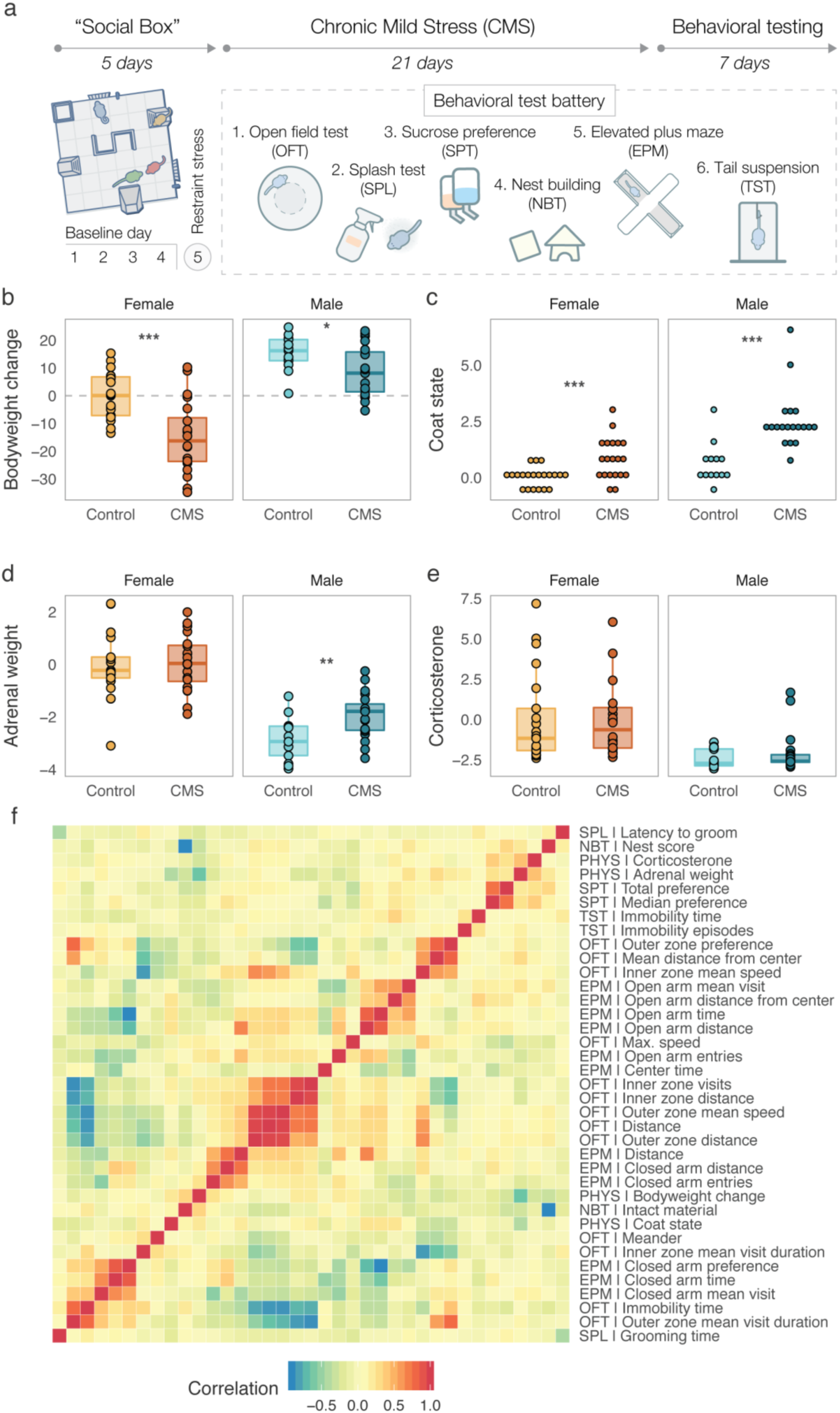
Outcomes of chronic mild stress (CMS) in males and females. (a) Experimental timeline. All groups underwent five days of Social Box (SB) monitoring. This consisted of four days of baseline monitoring followed by a 15-min acute restraint stress for all individuals prior to being re-introduced into the SB for a final 12-h dark phase monitoring period (day 5). After the SB, groups received three weeks of either control treatment (bodyweight and fur quality assessments two times a week) or chronic mild stress (CMS, see Methods for details). The following week, all groups underwent a behavioral test battery in the order depicted. (b) Batch-adjusted bodyweight change following three weeks of CMS. Both male and female CMS mice showed significantly reduced weight compared to controls. (c) Batch-adjusted coat state scores (higher means poorer fur quality) following CMS. Male and female CMS groups showed significant deterioration of their coat. (d) Batch- and initial bodyweight-adjusted adrenal weights. CMS increased adrenal size in males, but not in females. (e) Batch-adjusted plasma corticosterone level at sacrifice. No difference was observed between conditions. (f) Correlation structure of behavioral battery and physiological outcome data (37 readouts) across all samples (n = 73). (Boxplots: line – median, box limits – 1^st^ and 3^rd^ quartile, whiskers – 1.5 x IQR. Data is presented relative to female controls. Number of mice per condition: *Female*_*Control*_ = 21, *Female*_*CMS*_ = 20, *Male*_*Control*_= 13, *Male*_*CMS*_ = 19. \**p* < .05, \*\**p* < .01, \*\*\**p* < .001).

As expected, we found that both bodyweight change and cumulative coat quality were significantly reduced following CMS in both males and females (**Figure 2b-c**, Bodyweight: *F*(1, 69) = 75.5, *p* = 11×10^−13^, Coat quality: KW test, χ^2^(1) = 25.07, *p* = 6.5×10^−6^). Bodyweight-adjusted adrenal weights were increased after CMS in males only (**Figure 2d**, 2-way ANOVA, trend for a sex x condition interaction, *F*(1, 69) = 2.97, *p* = .089, followed by pairwise within-sex 2-sided t-tests: males: *t*(26.8) = −2.98, *p* = .006; Females: *t*(38.7) = -.17, *p* = .86). We did not observe an effect of CMS on plasma corticosterone levels at the time of sacrifice (**Figure 2e**, KW test: *χ*^2^(1) = .01, *p* = .9).

For all further analyses, these physiological outcomes were combined with the behavioral ones in a single dataset (the correlation structure of all the variables is depicted in **Figure 2f**). As expected, we observed that locomotion-related parameters such as distances in the OFT and in the EPM were related. Similarly, anxiety-dependent phenotypes such as open arm exploration in the EPM and distance from the center in the OFT showed a similar tendency. From these observations we can infer that our behavioral tests show the expected structure, clustering readouts classically associated to similar phenotypes.

### Sexually divergent effects of dominance on CMS outcomes

To explore how exposure to chronic stress shapes behavior in groups of mice, we investigated the major drivers of variance in the dataset containing all behavioral and physiological readouts following CMS (**Figure 2f**) using principle components analysis (PCA, **Figure 3a-d**). The first principal component (PC1), explained approximately 21% of the variance in the outcome data (**Figure 3a**). To our surprise, neither sex nor condition (CMS vs controls) appeared to capture variance contained in PC1 (**Figure 3b**, condition effect - *F*(1, 69) = .4, *p* = .52), but instead they were associated with PC2 and PC3 respectively (Supplementary **Figure 3**). Since none of the expected variables (sex, condition, or their interaction) contributed to PC1, the main source of variance in the dataset, we investigated whether social dominance was a contributing factor. We therefore tested the association between PC1 scores and DS (**Figure 3c**). Remarkably, baseline DS significantly predicted scores on PC1 in CMS individuals only and this association was in opposite directions between the two sexes (sex x DS interaction: *F*(1, 35) = 12.695, *p* = .0011). Thus, the principal source of variation in the outcome dataset could be described by an interaction between baseline dominance scores and sex in the CMS mice.

**Figure 3.**
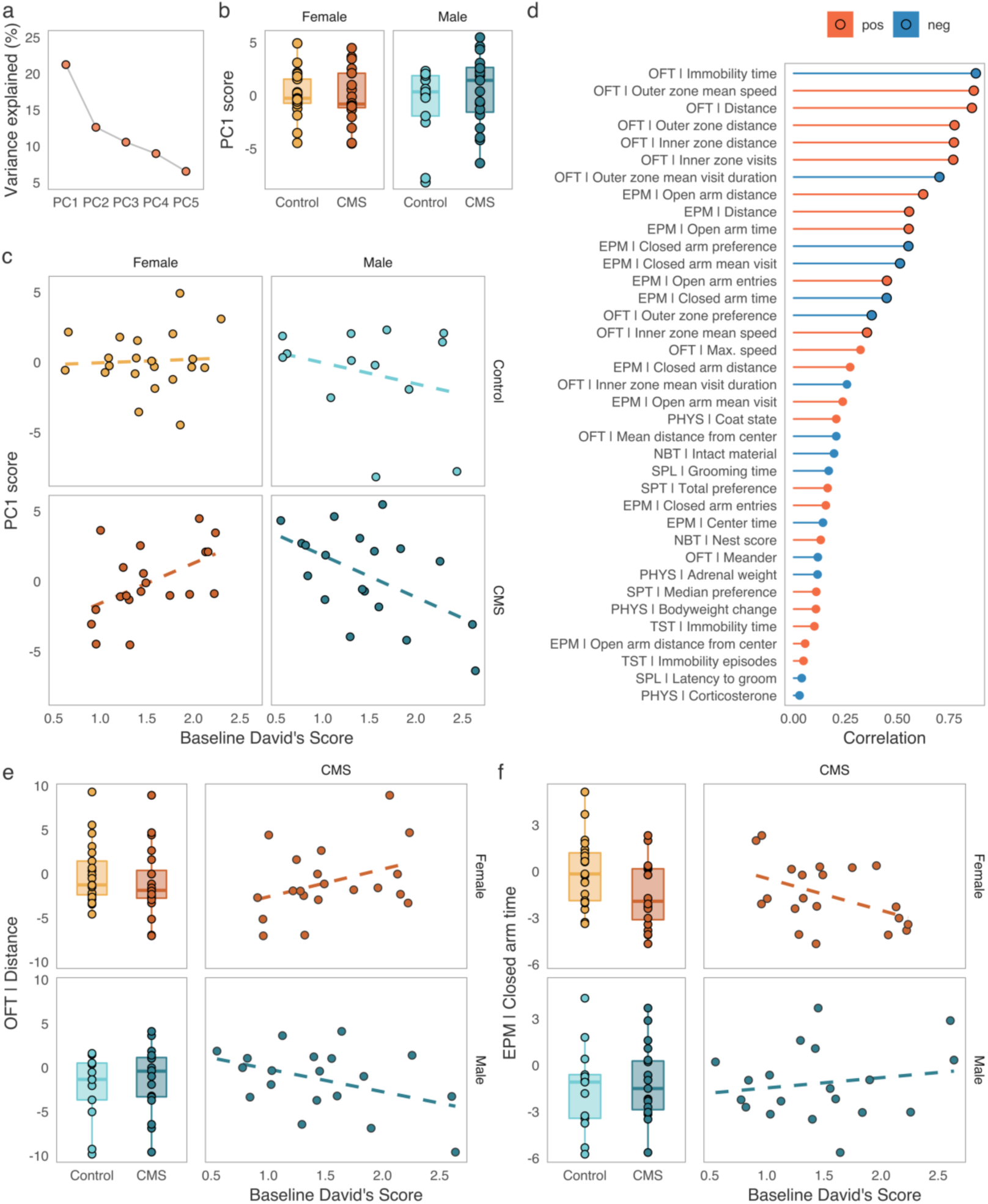
Opposing effects of baseline social dominance scores on behavioral outcomes following CMS. (a) Proportion of variance explained by the first five components of a principle components analysis conducted on the batch-adjusted behavioral and physiological outcome data. PC1 explains ca. 21% of the variance in this dataset. (b) PC1 is not significantly different between sexes or conditions, indicating that this component did not capture variance associated with either variable. (c) Association between baseline David’s Scores and PC1 in control and CMS individuals. Baseline dominance predicted scores on PC1, the major source of variance outcome data, in a sex-specific manner in the CMS group, but not in the control group. (d) Spearman’s rank correlations between PC1 and the physiological and behavioral outcome variables. The strongest associations for PC1 are variables derived from the open field test (OFT) and elevated platform maze (EPM). Black stroke around points identifies associations significant at *p* < .05 after adjustment for multiple testing (Bonferroni correction). (e-f) Examples of interactions between baseline dominance and sex on CMS behavioral outcomes. Males and females show significant opposite correlations between dominance and example of anxiety-like behavior. (Boxplots: line – median, box limits – 1^st^ and 3^rd^ quartile, whiskers – 1.5 x IQR. Scales for behavioral outcomes are relative to female controls).

To better understand the set of behaviors responsible for this association, we correlated PC1 scores with all the input features from the behavioral and physiological readouts (**Figure 3d**). We found that sixteen behaviors were significantly correlated with PC1 scores in this dataset (Spearman’s rank correlation, Bonferroni-adjusted *p* < .05). Among the strongest correlates of PC1 were measures derived from the OFT and EPM, and specifically features related to locomotion and anxiety-like behavior, such as distance traveled and visits to the anxiogenic regions of test chambers. Interestingly, these behaviors do not typically differentiate CMS and control individuals. Instead, CMS exposure created relationships between dominance and the outcome variables that were not present in controls (two examples are depicted in **Figure 3e-f**).

Finally, while the continuous nature of the of dominance provided by the DS affords power to assess correlations, social dominance hierarchies are typically thought of as ordinal. Since we had observed that the highest and lowest-ranking positions in the social hierarchy appeared most stable over time (**Figure 1c**), we ranked all individuals as belonging to one of three categories: *α -* dominant, *δ –* subordinate, or *β*/*γ* – neither dominant nor subordinate. We then used the scores on PC1 to better understand if any particular rank might be driving the differences observed in either sex following CMS. We found that *α*-males showed a trend toward a difference from the *β*/γ-ranking individuals on PC1 (*F*(1, 13) = 3.99, *p* = .067) and *δ*-males appeared to differ from the intermediate ranks (*F*(1, 13) = 4.85, *p* = .046) (**Figure 4a**). In females, only the *α*-ranking individuals seemed to differ from the intermediate ranks (*F*(1, 13) = 5.76, *p* = .032), while *δ*-females showed no differences (*F*(1, 12) = .002, *p* = .96) (**Figure 4a**). This suggests that while both dominance and subordination in male individuals may contribute to opposite behavioral responses to CMS, in females these effects are more likely to be driven by the dominant individuals only, which is consistent with our findings on rank stability.

**Figure 4.**
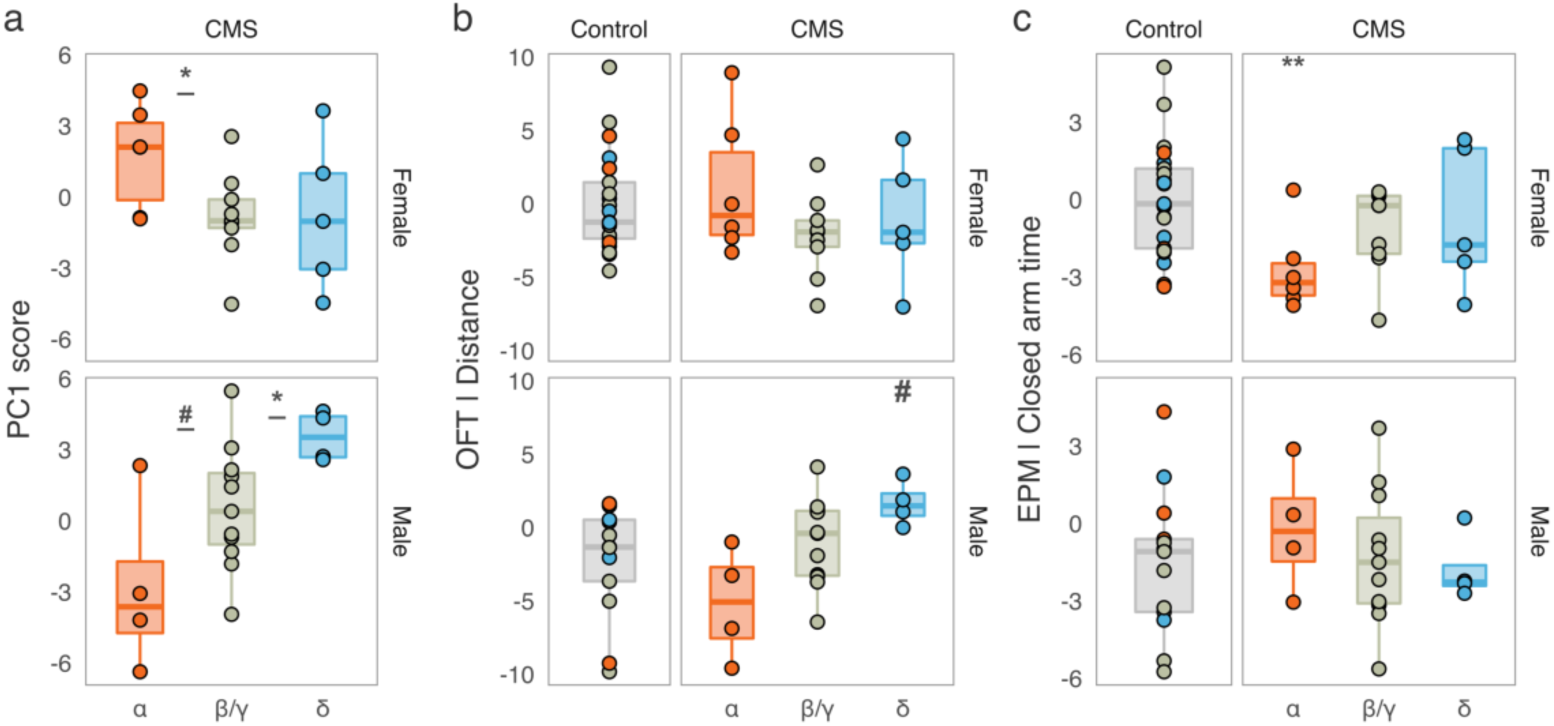
Rank-specific behavioral associations. (a) PC1 scores of CMS individuals show an effect of rank-belonging for *α*-females and δ-males and a trend for *α*-males, suggesting that specifically these ranks may be important drivers of the dominance x sex interaction. (b) Distance travelled in the OFT is an example of a behavioral readout for which CMS *δ*-males show a trend toward a difference from control males. (c) Time spent in the closed arms of the EPM is an example of a behavioral readout for which CMS *α*-females show a difference from control females (for full list, *see* Supplementary Figure 4). (Boxplots: line – median, box limits – 1st and 3rd quartile, whiskers – 1.5 x IQR. #p < .1, *p < .05, **p < .01. Scales for behavioral outcomes are relative to female controls).

To further clarify how the dominant and subordinate positions in the social hierarchy are related to CMS-specific behavior with respect to unstressed controls, we tested whether CMS *α*- & *δ*-males and CMS *α*-females differed from their sex-matched non-CMS controls on the 16 behavioral readouts that had shown significant associations with PC1. While *α*-males showed no significant differences from controls, CMS *δ*-males and CMS *α*-females each differed significantly from their controls on 4 out of 16 behaviors tested (nominal *p*-values, examples in **Figure 4b-c**, all values in Supplementary **Figure 4**). In particular, *δ*-males showed increased activity and reduced anxiety-like behavior after CMS exposure. Similarly, CMS *α*-females differed from controls on parameters associated with anxiety-like behaviors, such as preference between anxiolytic and anxiogenic zones of the EPM. To conclude, we were able to narrow down a portion of the dominance x sex interaction following CMS to the contribution of specific ranks, with CMS *δ*-males showing apparent increases in measures of overall activity (distance/speed in the OFT and EPM) and CMS *δ*-males and *α*-females showing an apparent reduction in anxiety-like behavior. Thus, we were able to identify a novel role for social rank belonging in modulating behavior after chronic stress in a sexually dimorphic way.

## Discussion

Social behavior in general and social dominance in particular are important contributors to individual differences(24). As such, they may also shape how individuals respond to environmental challenges. Here we have confirmed that both male and female socially housed mice establish social dominance hierarchies, which are relatively stable over time and resistant to acute perturbations. In our hands, female hierarchies were less despotic and had lower directional consistency, as has been previously observed(19), suggesting that females may be maintaining a less rigid structure than males. This is supported by our finding that only the top rank females showed significant stability over time, whereas in males both the most subordinate and the most dominant ranks were appeared stable. Further research is needed to ascertain if this finding is confined to our paradigm and specific set of measurements, or if it represents a true sex difference in social dominance hierarchies. Additionally, our data suggest that an individual’s position in the hierarchy carries different implications for overall behavior in each sex. We observed that for groups living in the enriched environment of the SB apparatus, dominance in males but not in females was associated with overall locomotion, proportion of time spent outside the nest, and exploration entropy. These associations likely reflect territorial or patrolling behavior in males, which may be less relevant to female social hierarchies.

As hypothesized, occupancy of different positions in the social hierarchy conferred varying levels of responsiveness to the challenges posed by chronic mild stress. Previous investigations have rarely found associations between social dominance and response to chronic stress(29). An important exception is a recent study by Larrieu et al.(30) in groups of male mice exposed chronic social defeat. The authors found increased susceptibility to chronic social defeat for dominant males, but in contrast to our results, no real behavioral alterations for subordinates. It is important to note, however, that chronic social defeat and CMS are profoundly different paradigms. Social defeat is strongly tied to social dominance and might be perceived as loss of status more than other stressors(29). Our use of CMS allowed us to investigate both sexes under comparable levels of stress. Nevertheless, both the study from Larrieu and colleagues and ours highlight that social status can influence an individual’s response to long-term life events.

Crucially, in our hands, the effects of preexisting dominance on stress outcomes were sexually divergent, such that the association between dominance and anxiety-like and locomotor behavior following CMS was in opposite directions between males and females. In particular, subordinate males appeared to display hyperlocomotion, while dominant females displayed increased boldness (reduced anxiety-like behavior) compared to non-CMS controls. Overall, our data suggest that an individual’s position within a social structure can influence their behavioral response to chronic stress in a sex-specific fashion.

Altogether our findings suggest an intriguing possibility. Given that male social hierarchies are likely antagonistic, we speculate that social living carries an especially high cost for all subordinate males, who are the recipients of most antagonistic interactions. Conversely, female hierarchies may contribute to more affiliative social interactions, and thus social context may carry a net benefit for females, with the highest benefit gained by the dominant females. We speculate that this positioning as the most advantaged and disadvantaged individuals may confer higher behavioral flexibility and results in the strongest behavioral change upon exposure to environmental challenges.

Since we decided to maintain social context throughout our experimental design, the current work did not allow for the assessment of the effect of group-versus single-housing on CMS outcomes. Given this constraint, we were not able to confidently assess the difference in how CMS was experienced by each sex in groups as opposed to if they had been single-housed. However, since we were interested in the prediction from baseline dominance, we did not wish to remove the salience and thereby the effect of social context.

Additionally, while CMS produced some of the expected physiological changes (i.e., reduction of bodyweight gain, reduced coat quality, adrenal weight increase), we did not observe several of the behavioral phenotypes often found using similar protocols (e.g., hyperlocomotion, anhedonia, passive coping(31)). While we have sufficient evidence that CMS individuals experienced significant amounts of stress, we are not able to determine if the absence of some of these behavioral signatures of CMS is a result of the maintenance of social context throughout the protocol or if it should be attributed to other unknown factors. We are, however, not the first to observe no increase in adrenal size or no decrease in sucrose preference in female CD1 mice(32). Additionally, we did not observe any changes in basal corticosterone levels. This may be due to the fact that our blood sampling was performed one week after the end of the CMS paradigm, which may have been sufficiently long for corticosterone levels to return to normal.

Finally, we employed the David’s Score as a continuous linear indicator of social dominance for the additional statistical power that this approach provides. Dominance hierarchies, however, are more commonly thought of as ordinal, and we lacked sufficient sample sizes per rank and condition to be able to reliably quantify the contribution of each rank to the behavioral outcomes of chronic stress. Further research is needed to replicate and extend these findings to specific social ranks.

While were not able to provide a direct comparison between single- and group-housed animals, our data suggest that the existence of a social hierarchy in groups of mice might contribute to increased variability in behavioral outcomes after chronic treatment generating rank-specific responses. Moreover, this effect could be especially relevant when studying sex differences. Often, housing conditions (single vs. group) are not taken into consideration as a variable of interest. Based on the findings reported here, we speculate that housing conditions might have contributed to discordant behavioral findings in studies of stress and sex(31). Our results argue for considering group-derived individual differences and, in particular, dominance status, in the design of experiments, especially when investigating the contribution of sex differences to stress response.

Taken together, this work suggests that social dominance might influence the perception of and reaction to chronic stress differently for male and female mice. While there has been some work looking into the effects of dominance on stress susceptibility in males(30), very little is known about female social dominance and its contribution to stress coping. Our work emphasizes both the need for exploring the stress response in the presence of conspecifics in a more naturalistic manner, and the importance of recognizing that the same social factors may carry divergent consequences for the behavior of males and females.

## Materials and Methods

### Animal housing and care

Male and female ICR CD-1 mice at 7-9 months old were employed for all experiments (Charles River, Sulzfeld, Germany). Mice were housed in groups of four in the animal facilities of the Max Planck Institute of Psychiatry in Munich, Germany, from weaning and were maintained under standard conditions (12L:12D light cycle, lights on at 07:00 AM, temperature 23 ± 2°C) with food and water available *ad libitum*. All experiments were approved by and conducted in accordance with the regulations of the local Animal Care and Use Committee (Government of Upper Bavaria, Munich, Germany).

### Behavior in a semi-naturalistic environment

#### The “Social Box” paradigm

The “Social Box” is a behavioral arena wherein groups of mice live under continuous observation over a period of several days(23,24). Mouse identities were maintained using fur markings in four different colors(24).

The entire observation period was recorded using cameras mounted above each arena. Videos were then compressed and analyzed using a custom-made automated tracking system which determines mouse locations over time(23). From the location data we inferred agonistic interactions and a variety of other behavioral readouts as described previously(24).

#### Social dominance

Social dominance was assessed using the David’s Score (DS), a measure based on the pairwise directionalities and numbers of agonistic interactions in a group(26). The DS was normalized to group number (*n* = 4), creating a continuous range between 0 (least dominant) and 3 (most dominant). The steepness of the social hierarchy was characterized as described ref. (33) by using the slope of a line fitted to the DS from a ranked DS using Ordinary Least Squares regression. We used an implementation of this procedure made available in the “steepness” R package.(34) Despotism was defined as the fraction of the group’s total number of chases that were initiated by the highest-ranking individual. Directional consistency represents the average pairwise fraction of social interactions that occur in the direction from the individual who displayed more instances of an agonistic behavior to the individual who displayed fewer instances(35,36). Finally, we used Landau’s modified *h*’ to assess the linearity of a social hierarchy, as described in ref.(28). We calculated both directional consistency and Landau’s modified *h*’ using functions made available in the R package “compete”(37).

#### Acute restraint

Before the beginning of the fifth night in the Social Box, mice were removed from the SB and restrained in a ventilated tube for 15 minutes. To account for the smaller size females, we employed a smaller sized ventilated tube, to ensure the same degree of movement restriction between sexes. At the end of the stressors, groups of mice were put back in their original SB and tracked for an additional 12 hours.

### Chronic mild stress protocol

Two separate batches of mice were exposed to three weeks of chronic mild stress prior to the behavioral test battery. A random combination of two stressors per day (one in the a.m. and one in the p.m. hours) was chosen among the followings: acute restraint in the dark (15 min), acute restraint in bright light (15 min, ∼200 lux), acute restraint witnessing (half of the group at a time was restrained and placed inside the cage, 15 min each), removal of nesting material (24 h), cage-tilt 30° along the vertical axis (6 h), no bedding or nesting material (8 h), wet bedding (6 h), water avoidance (15 min), cage change (fresh cage every 30 min for a total of 4 h), cage switching (mice are assigned the cage of another group of the same sex), overcrowding (8 mice per cage, 1 h). For the water avoidance stress, an empty rat cage (395 × 346 cm) was filled with room temperature water. Mice were placed on a platform (10 × 12 cm), 2 cm above the water level, for 15 minutes.

On days 1, 3, 7, 10, 14, 17 and 21 both CMS and control mice were weighed. During the weighing session, their coat state was scored on a scale 0 to 3 according to the following criteria:

0) Bright and well-groomed coat. Clean eyes. No wounds.
1) Less shiny and less groomed coat OR unclean eyes. No wounds.
2) Dirty and dull coat AND/OR small wounds and not clear eyes.
3) Extensive piloerection OR alopecia with crusted eyes OR extensive wounds.

Cumulative coat state was calculated as the sum of the seven daily scores.

### Behavioral battery

The day after the last stressor, mice started a behavioral test battery consisting of the open field test (OFT), two-hour sucrose preference test (SPT), grouped sucrose preference test, the splash test (SPL), the nest building test (NBT), the elevated plus maze (EPM), a grouped sucrose preference, and the tail suspension test (TST). Throughout the testing period, mice were maintained in their original groups and habituated to the testing room for at least one hour prior the start of the test. Forty-eight hours after the last test, mice were terminally anesthetized in isoflurane and sacrificed. Terminal bodyweight, plasma, adrenal glands and thymus were collected. Adrenal glands and thymus were cleaned from fat tissue and weighed. Absolute values were adjusted to bodyweight using the bodyweights collected on day 1.

#### Open field test

On the day following the last stressors (day 22), mice locomotor activity and exploratory behavior were assessed in the open field test for 10 min. The apparatus consisted in round arenas (diameter 38 cm) made of black polyvinylchloride (PVC) under dim illumination (3 lux). Mice were automatically tracked with ANYmaze Video Tracking System 6.13 (Stoelting, IL, USA). The space was virtually divided in an inner zone (diameter 16 cm) and an outer zone. Total distance traveled, distance from the center, speed, and turn angle were calculated across the full 10 minutes. In addition, distance traveled, speed, visits, and time spent in each of the subdivisions were used as parameters.

#### Two-hour daily sucrose preference test

Twenty-four hours after the OFT, the anhedonia phenotype was tested with a modified version of the sucrose preference test. Each group was assigned a test cage containing one water bottle and one bottle with 2% sucrose. One mouse per group at a time was placed in the test cage for two hours, across three consecutive days during the light phase (days 23, 24, and 25). At the end of each session, the bottles were weighed. At the end of the test the amounts of water and sucrose consumed were summed across the three sessions. Sucrose preference was calculated as 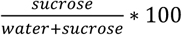.

#### Grouped sucrose preference test

On day 27, sucrose preference was tested at a group level. Each group was given a bottle of water and a bottle of 2% sucrose within their home-cage. Their sucrose preference was calculated after 24 hours as above. A grouped sucrose preference value was obtained for each group.

#### Splash test

On day 24, during the dark period, mice were tested in the splash test under dim light (3 lux). Mice were placed in their test cage for 5 minutes prior being sprayed on their dorsal coat twice (approximately 1 ml) with 10% sucrose solution. Mice were recorded for 5 minutes and total time spent grooming, and latency to the first grooming bout was manually scored using Solomon Coder 17.03.32 (https://solomon.andraspeter.com/) by an experimenter blind to sex and condition.

#### Nest building test

During the third day of the two-hour sucrose preference, mice in the test cage were given a small square cotton pad of approximately 23 g. The cotton pad was weighed at the beginning of the test and at the end of the two hours and the percentage of intact material was calculated. The built nest was scored from 0 to 4 according to the following criteria:

0) Material untouched
1) Material partially torn (50-90% remaining intact)
2) Material mostly shredded but often no identifiable nest site/scattered around
3) Material accumulated in an identifiable nest site, but the nest is flat
4) A (near) perfect nest: material fine shredded, doughnut like with walls higher than the mouse

For nests matching only partially the description (i.e., identifiable flat nest, but less than 50% of torn material), half points were assigned.

#### Elevated plus maze

On day 26, during the light phase, anxiety phenotype was assessed using the elevated plus maze test. An apparatus composed of four arms made of grey polyvinylchloride (PVC), two open without walls, two enclosed by 14 cm walls and a central platform (5 × 5 cm). The apparatus was placed 33 cm from the ground under dim illumination (3 lux). Mice were placed on the central platform facing the open arms and let free to explore the apparatus for 10 min. Mice were automatically tracked using ANYmaze Video Tracking System 6.13 (Stoelting, IL, USA). Number of entries in each arms, time, and distance were calculated. In addition, closed arm preference was calculated as 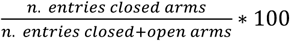.

#### Tail suspension test

Stress coping behavior was assessed using the tail suspension test on day 28. Mice were hung by their tail 50 cm above a surface and their behavior recorded for 6 min. Immobility was automatically scored using ANYmaze Video Tracking System 6.13 (Stoelting, IL, USA) and number of immobility episodes and total time immobile were used as parameters.

### Corticosterone assessment

At sacrifice, trunk blood was collected in EDTA-coated tubes. Blood was centrifuged at 1,000g for 15 min at 4°C. Plasma was retrieved and corticosterone levels were measured using [125I] radioimmunoassay kit (MP Biomedicals), according to the manufacturer’s instructions.

### Statistical analyses

All statistical analyses were performed in R version 3.5.0(38). The tests employed for each specific analysis are reported in the Results section. Outcome data distributions were tested for deviations from normality (Shapiro-Wilk test) and heteroscedasticity (Levene’s test). Whenever normality was violated and the data could not be transformed to fit a normal distribution, non-parametric tests were employed. Violations of homogeneity of variances are reported with each test. For the special case reported in Supplementary **Figure 4**, given the pronounced inequality in sample sizes, we used permutation-based analyses of variance, as implemented in the lmPerm R package(39). The effect of experimental batch on the behavioral outcome data following CMS was adjusted for by using the standardized residuals of a linear model with each variable of interest as outcome and batch as a factorial predictor. Principle components analysis was performed using singular value decomposition on scaled and centered data.

## Abbreviations

CMS: chronic mild stress;
SB: Social Box;
DS: David’s Score;
OFT: open field test;
EPM: elevated plus maze;
PCA: principle components analysis.

## Acknowledgments

We thank Yair Shemesh, Oren Forkosh, Markus Nussbaumer, and Chadi Touma for their assistance in setting up the “Social Box” paradigm. Thanks to Jessica Keverne for English writing support and advice. A.C. is the incumbent of the Vera and John Schwartz Family Professorial Chair in Neurobiology at the Weizmann Institute and the head of the Max Planck Society–Weizmann Institute of Science Laboratory for Experimental Neuropsychiatry and Behavioral Neurogenetics. This work is supported by: an FP7 Grant from the European Research Council (260463, A.C.); a research grant from the Israel Science Foundation (1565/15, A.C.); the ERANET Program, supported by the Chief Scientist Office of the Israeli Ministry of Health (3-11389, A.C.); the project was funded by the Federal Ministry of Education and Research under the funding code 01KU1501A (A.C.); I-CORE Program of the Planning and Budgeting Committee and The Israel Science Foundation (grant no. 1916/12 to A.C.); Ruhman Family Laboratory for Research in the Neurobiology of Stress (A.C.); research support from Bruno and Simone Licht; the Perlman Family Foundation, founded by Louis L. and Anita M. Perlman (A.C.); the Adelis Foundation (A.C.); and Sonia T. Marschak (A.C.). S.K. and E.B. are supported by the International Max Planck School for Translational Psychiatry (IMPRS-TP).

## Author contributions

Conceptualization, SK, EB, and AC; Methodology, SK, EB, MS; Data analysis, SK, EB, RS, CF; Writing, Review & Editing, SK, EB, MVS, and AC; Funding Acquisition, AC; Supervision, AC.

## Competing interests

The authors declare no competing interests.

## Data availability

All data used to support the findings of this work are available from the corresponding author upon reasonable request.

## Code availability

The code used in performing the analyses and producing the Figures for this manuscript is freely accessible in a GitHub repository https://stoyokaramihalev.github.io/CMS_Dominance/.

The MATLAB-based mouse tracking system is available from the corresponding author upon request.

## Supplementary Information

**Supplementary Figure 1.**
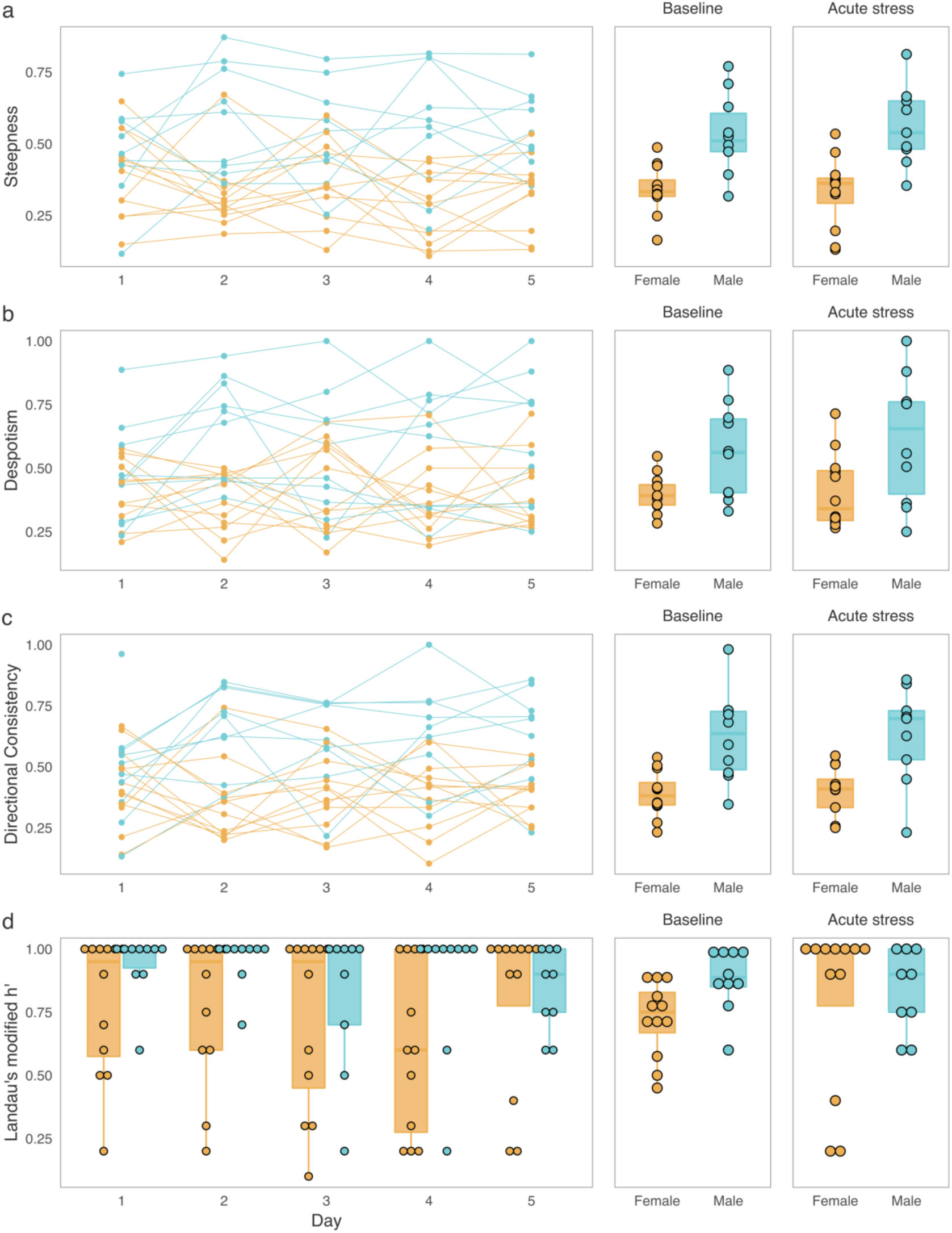
Social hierarchy properties in male and female groups. **(a-d)** The following properties of social hierarchies were computed for each group of mice: steepness, despotism, directional consistency, and Landau’s modified h’ – a measure of the linearity of a social hierarchy. Male groups had overall higher for steepness (*F*(1, 19) = 21.18, *p* = 1.9×10^−4^), despotism (*F*(1, 20) = 7.83, *p* = .011, *note:* variances were heterogenous between conditions), and directional consistency (*F*(1, 19) = 24.4, *p* = 9×10^−5^). Means of Landau’s modified h’ over the baseline days also differed between the sexes (KW test: χ^2^(1) = .5.84, *p* = .016). None of the measures showed evidence of a sex x acute stress interaction.

**Supplementary Figure 2.**
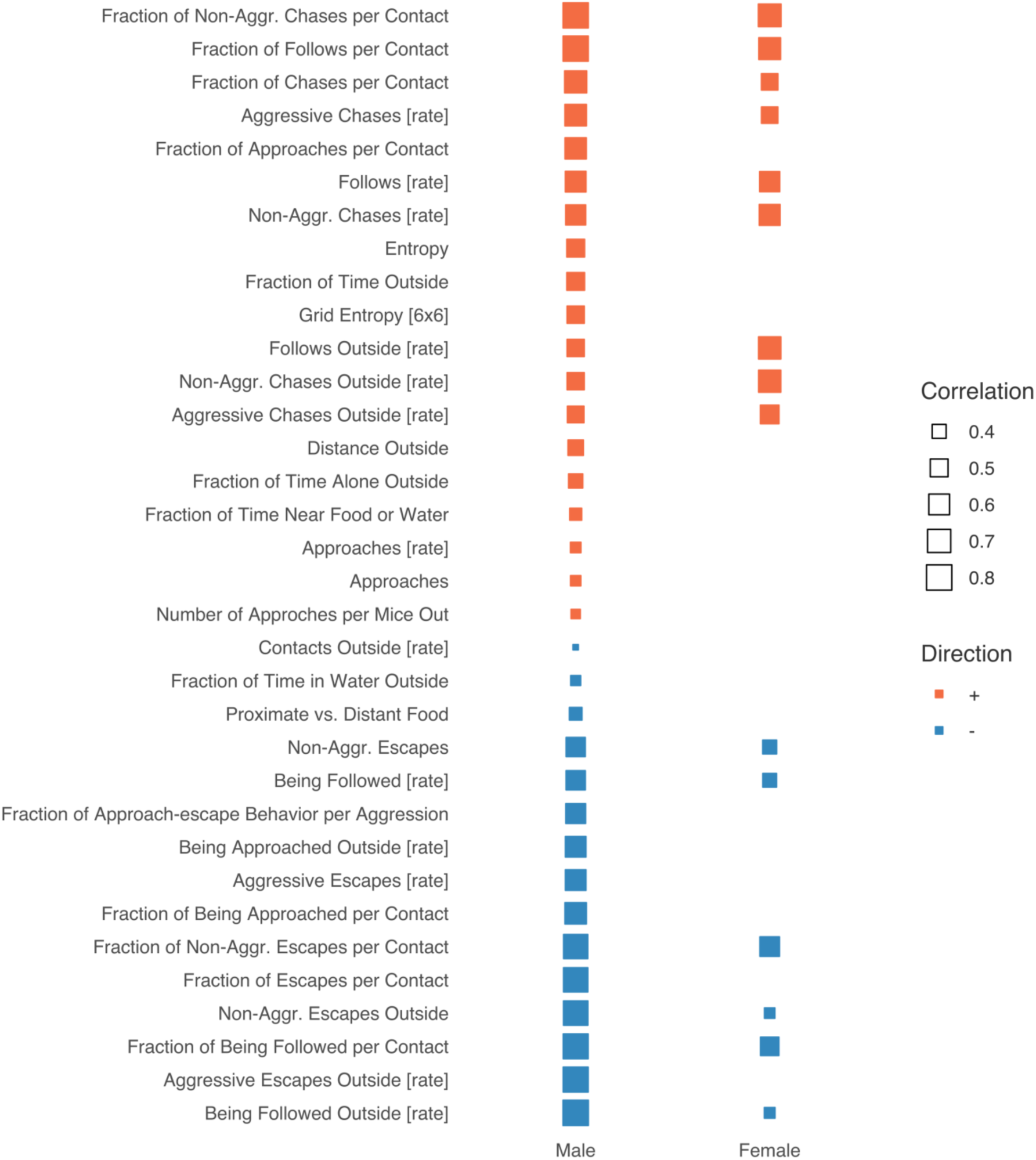
Associations between David’s scores and other behaviors measured in the Social Box. Depicted are cumulative baseline David’s scores correlations with batch-adjusted measures from the Social Box (averages of each behavioral readout for Social Box days 2-4). Only behaviors that are not used for building the DS (34 out of 36) and only correlations with a q-value < 0.1 following multiple testing correction are displayed (Benjamini-Hochberg method, for a full list of the behavioral readouts tested here, *see* Methods). David’s Scores show relatively similar correlation patterns with other behaviors between the sexes, indicating that social dominance status may have largely similar effects on overall behavior for each sex. Nevertheless, several behavioral readouts, including Distance Outside, Entropy / Grid Entropy [6×6], and Fraction of Time Outside show associations with David’s scores in males that are conspicuously absent in females (**Figure 1c**).

**Supplementary Figure 3.**
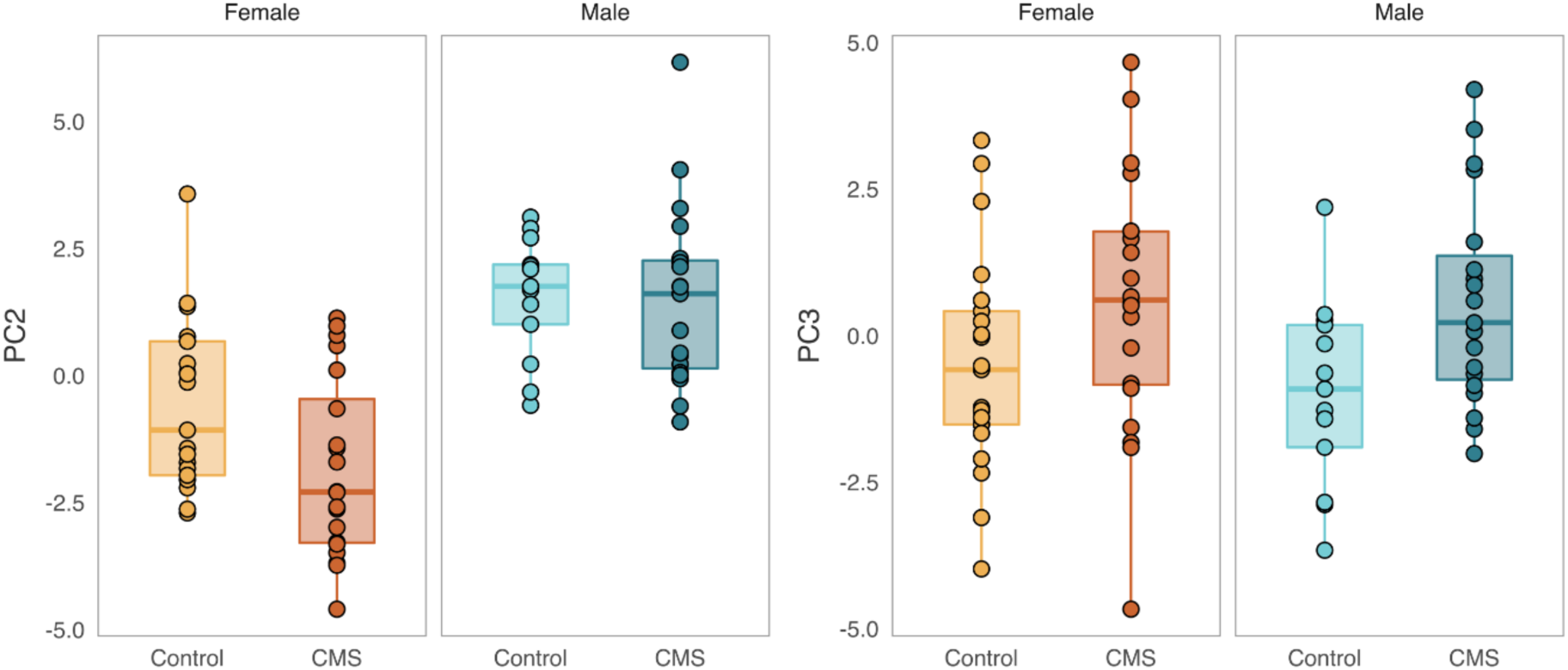
PC2 and PC3 capture variance associated with sex and condition. Depicted are scores for two further principle components from the analysis described in **Figure 3**. Individual scores on PC2 (12.6% variance explained) differ between the sexes (*F*(1, 69) = 51.5, *p* = 6.4×10^−10^), while PC3 (10.6% variance explained) captures an effect of chronic mild stress (CMS, *F*(1, 69) = 7.89, *p* = .006).

**Supplementary Figure 4.**
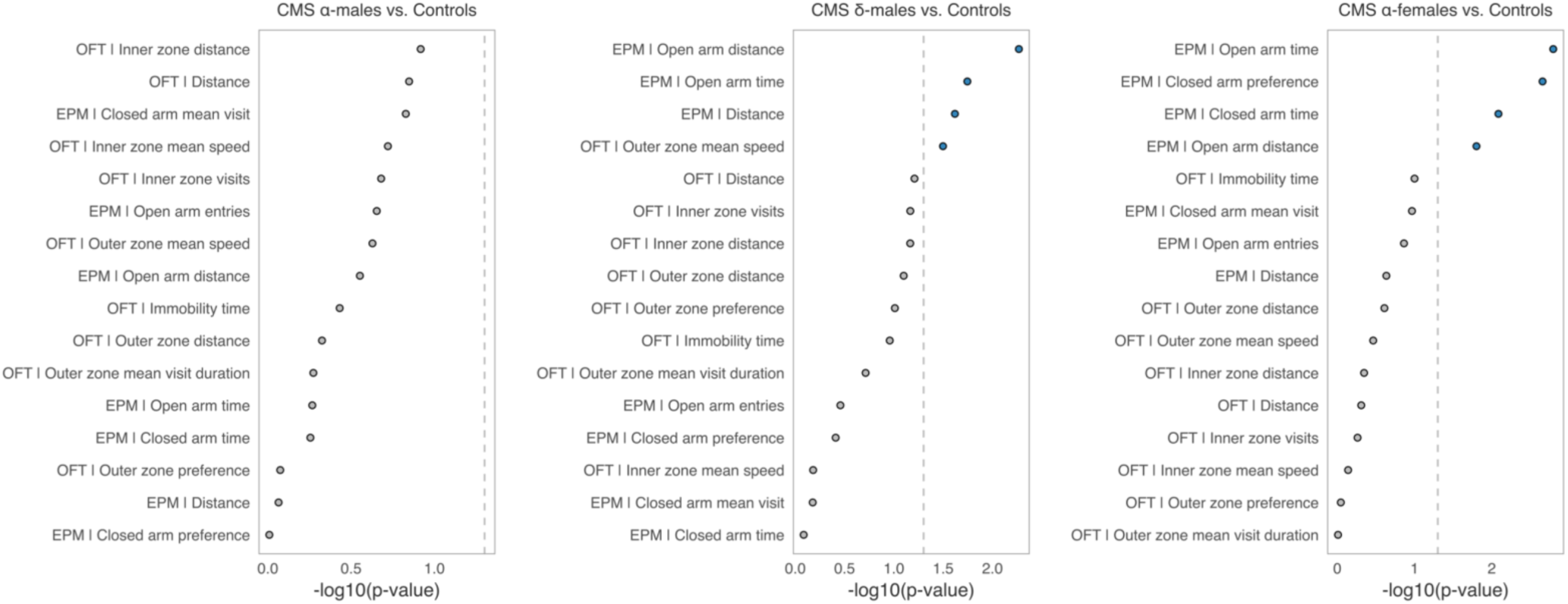
CMS *α*-females, CMS *α*-males, and CMS *δ*-males, compared to the sex-matched control groups. Depicted are the negative log10 *p-*values for each comparison between CMS *α-* & *δ*-males vs. all male controls, as well as CMS *α*-females versus all female controls (permutation-based analysis of variance; colored points represent nominally significant comparisons at *p* < .05).

## Notes

https://stoyokaramihalev.github.io/CMS_Dominance/

